# Local membrane charge regulates ல2 adrenergic receptor coupling to Gi

**DOI:** 10.1101/400408

**Authors:** M.J. Strohman, S. Maeda, D. Hilger, M. Masureel, Y. Du, B.K. Kobilka

## Abstract

G protein coupled receptors (GPCRs) are transmembrane receptors that signal through heterotrimeric G proteins. Lipid modifications anchor G proteins to the plasma membrane; however, little is known about the effect of phospholipid composition on GPCR-G protein coupling. The β_2_ adrenergic receptor (β_2_AR) signals through both G_s_ and G_i_ in cardiac myocytes where studies suggest that G_i_ signaling may be cardioprotective. However, G_i_ coupling is much less efficient than G_s_ coupling in most cell-based and biochemical assays, making it difficult to study β_2_AR-G_i_ interactions. To investigate the role of phospholipid composition on G_s_ and G_i_ coupling, we reconstituted β_2_AR in detergent/lipid mixed micelles and found that negatively charged phospholipids (PS and PG) inhibit β_2_AR-G_i3_ coupling. Replacing negatively charged lipids with neutral lipids (PC or PE) facilitated the formation of a functional β_2_AR-G_i3_ interaction that activated G_i3_. Ca^2+^, known to interact with negatively charged PS, facilitated β_2_AR-G_i3_ interaction in PS. Mutational analysis suggested that Ca^2+^ interacts with the negatively charged EDGE motif on the carboxyl-terminal end of the αN helix of G_i3_ and coordinates an EDGE-PS interaction. These results were confirmed in β_2_AR reconstituted into nanodisc phospholipid bilayers. β_2_AR-G_i3_ interaction was favored in neutral lipids (PE and PC) over negatively charged lipids (PG and PS). In contrast, basal β_2_AR-G_s_ interaction was favored in negatively charged lipids over neutral lipids. In negatively-charged lipids, Ca^2+^ and Mg^2+^ facilitated β_2_AR-G_i3_ interaction. Taken together, our observations suggest that local membrane charge modulates the interaction between β_2_AR and competing G protein subtypes.

## Introduction

A third of all FDA approved pharmaceutical drugs function by modulating the activity of G protein coupled receptors (GPCRs)^1^, a large receptor superfamily. GPCRs catalyze the activation of heterotrimeric G proteins, which in turn initiate a multitude of signaling cascades that alter cellular function.

G proteins are also a large superfamily, grouped into 4 subfamilies (G_s_, G_i/o_, G_q/11_, G_12/13_) encoded by 16 different genes^2^. Each subfamily activates distinct signaling pathways,and functional effects are cell-type specific. Most GPCRs can signal through more than one G protein subfamily, and ongoing research attempts to identify mechanisms that regulate G protein selectivity within a cell^2^.

Here we investigate the dual G protein selectivity of the β_2_ adrenergic receptor (β_2_AR), a prototypical GPCR that mediates the fight-or-flight response. The dual G protein selectivity of β_2_AR is best characterized in heart muscle (cardiac myocytes) where activation of G_s_ increases contraction rate and activation of G_i_ decreases it. β_2_AR activity is stimulated by the hormone epinephrine. In healthy neonatal cardiac myocytes, epinephrine stimulated β_2_AR immediately activates G_s_, but after 10-15 minutes β_2_AR signals predominantly through G_i_^3^. Of interest, G_i_ activation is impaired if β_2_AR internalization is blocked^4^. Also, G_i_ does not interact with a modified β_2_AR that internalizes but does not recycle to the plasma membrane^5^, and WT β_2_AR that internalizes but is pharmacologically blocked from recycling^6^. Taken together, these observations demonstrate that β_2_AR-G_i_ interaction is regulated spatially and temporally, and interaction occurs after epinephrine stimulated trafficking of β_2_AR.

β_2_AR-G_i_ signaling plays a complex role in heart disease. While its anti-apoptotic effects^7,8^ may prevent ischemic-reperfusion injury^9^, G_i_ activation comes at the price of decreased contractility, which is problematic in diseases where the heart is already sufficiently weakened, such as in heart failure^10^, Takotsubo syndrome^11,12^, and ischemia^13^. Notably, in heart failure G_i2_ is upregulated^14-16^, β_1_AR (strictly G_s_ coupled) is downregulated^17-20^, while G_s_^16^, G_i3_^16^, and β_2_AR^18^ expression levels are unchanged. These changes may increase β_2_AR-G_i2_ coupling relative to the β_2_AR-G_s_ coupling.

The mechanism that initiates β_2_AR-G_i_ signaling in the healthy heart is not fully understood. Multiple biochemical mechanisms may play a role. PKA phosphorylation of β_2_AR has been reported to increase G_i_ coupling *in vitro*^21^ and in HEK cells^22^, but in cardiac myocytes, β_2_AR-G_i_ coupling is PKA independent^3^. In addition, GRK2 phosphorylation of β_2_AR has been suggested to increase G_i_ coupling^23^, but other investigators have reported that dephosphorylation is critical for β_2_AR recycling to the plasma membrane, and β_2_AR-G_i_ interactions^6^. Therefore, we sought to discover mechanisms that modulate β_2_AR-G_i_ coupling.

The epinephrine-stimulated trafficking of β_2_AR (internalization and plasma membrane recycling) may influence the composition of phospholipids surrounding the β_2_AR. It has been shown that negatively charged phospholipids stabilize an active conformation of the β_2_AR and enhance its affinity for epinephrine^24^. Here we examine the effect of phospholipid charge on β_2_AR interactions with G_s_ and G_i_.

## Results

Epinephrine activates β_2_AR by stabilizing a conformation that is recognized by G protein. This conformation is partially stabilized by epinephrine and fully stabilized by the addition of G protein^25-27^. The conformational change can be detected using a modified β_2_AR labeled on Cys265 at the cytoplasmic end of TM6 with an environmentally sensitive fluorophore, monobromobimane (mB-β_2_AR, see methods)^28^. Epinephrine and G protein interaction red-shifts the emission maximum (*λ*max, the wavelength where fluorophore emission intensity is greatest) and decreases the intensity of mB-β_2_AR (Fig. 1a). Because *λ*max is independent of mB-β_2_AR concentration, it is a more reliable indicator of β_2_AR conformation than fluorescence intensity. We therefore monitored changes in *λ*max of mB-β_2_AR to detect G protein coupling.

**Figure 1.**
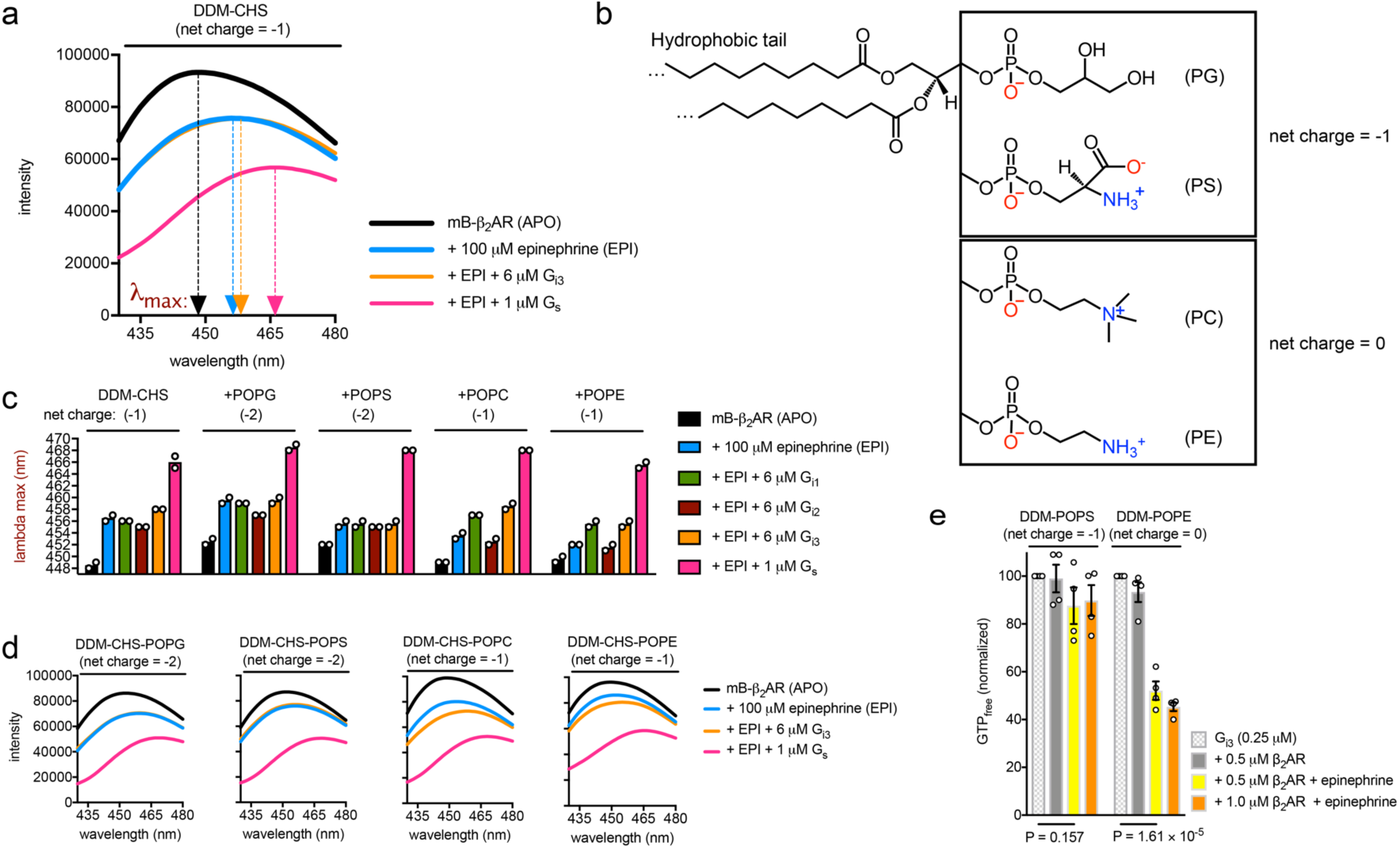
Effect of phospholipids on mB-β_2_AR-G protein interaction in DDM mixed micelles a. **a.**mB-β_2_AR emission spectra in the presence and absence of epinephrine and G protein (G_s_ and G_i3_) in DDM-CHS micelles (4:1 DDM:CHS). Arrows point to the lambda max value, i.e. the wavelength where mB emission intensity is greatest. **b.** Structure of phospholipids, categorized by their net charge: Phosphatidylglycerol (PG) and phosphatidylserine (PS) have a net charge of −1. Phosphatidylcholine (PC) and phosphatidylethanolamine (PE) have a net charge of 0 (i.e. net neutral). Negatively charged groups highlighted in red. Positively charged groups highlighted in blue. **c.** mB-β_2_AR interaction with epinephrine and G protein (G_i1_, G_i2_, G_i3_, and G_s_) as assessed by changes in mB lambda max. Interaction was assessed in DDM-CHS micelles (4:1 DDM:CHS) and in DDM-CHS micelles containing POPG, POPS, POPC, or POPE lipids (3:1:1 DDM:CHS:Lipid). **d.** Selected mB-β_2_AR emission spectra from panel C, showing spectra shifts induced by epinephrine, G_i3_, and G_s_. **a-d.** The indicated net charges are net molecular charge, not net micelle charge. CHS, POPG, and POPS each have a net charge of −1. DDM, POPC, and POPE are net neutral. mB-β_2_AR concentration is 300 nM. Data are mean of two independent experiments. **e.** GTP turnover in the presence and absence of β_2_AR (0.5 µM vs. 1 µM) and saturating epinephrine, in 4:1 DDM:POPS vs. 4:1 DDM:POPE. Data were normalized relative to turnover by G_i3_ (0.25 µM) in the absence of β_2_AR (shown in Supplementary figure 3). Data are mean +/- s.e.m of four independent experiments. Statistical significance was determined by using a two-sided Student’s *t*-test.

### Negatively charged phospholipids decrease β_2_AR coupling to G_i_

In the presence of epinephrine, we observe a change in intensity and *λ*max of mB-β_2_AR following the addition of G_s_ in a detergent mixture containing n-dodecyl-*β*-D- maltopyranoside (DDM) and cholesteryl hemisuccinate (CHS) that is commonly used for biochemical study of GPCR/G protein complexes (Fig. 1a). In contrast, the coupling efficiency of mB-β_2_AR-G_i3_ was relatively weak (Fig. 1a). Next, we compared the coupling efficiency of mB-β_2_AR-G_i_ in DDM-CHS mixtures with different phospholipids incorporated (Fig. 1b,c). While we were unable to detect interactions of mB-β_2_AR with G_i1_, G_i2_ or G_i3_ in the presence of negatively charged lipids (POPS and POPG), we observed a weak interaction with G_i1_ and G_i3_ in neutral lipids (POPE and POPC) (Fig. 1c,d). This result suggested that negatively charged lipids may repel G_i1_ and G_i3_ interaction with β_2_AR, despite the fact that negatively charged lipids enhance epinephrine binding affinity, as previously reported^24^. Given that mB-β_2_AR-G_i1_ and mB-β_2_AR-G_i3_ interaction appeared comparable, we narrowed our focus on mB-β_2_AR-G_i3_ interaction because G_i3_ (not G_i1_) is expressed in the heart^16^.

We also observed that negatively charged CHS decreased mB-β_2_AR-G_i3_ coupling (Supplementary Fig. 1). This effect was minimized in acidic buffers known to protonate (and neutralize) CHS^29^, supporting our hypothesis that negatively charged lipids decrease β_2_AR-G_i_ coupling. Given reports that PKA phosphorylation of β_2_AR increases β_2_AR-G_i_ interaction *in vitro*^21^, we also tested the effect of PKA phosphorylation, but no B-β_2_AR-G_i3_ enhancement was observed (Supplementary Fig. 2), suggesting that phosphorylation does not potentiate β_2_AR-G_i3_ interaction under our experimental conditions, and that other mechanisms may enhance β_2_AR-G_i3_ interaction.

In subsequent experiments, we omitted negatively charged CHS in order to assess the effect of phospholipid charge on mB-β_2_AR-G_i3_ interactions. We tested whether the increased mB-β_2_AR-G_i3_ interaction we observed in neutral lipid represented functional interaction. Indeed, β_2_AR stimulated GTP turnover was detected in DDM micelles containing POPE (net neutral lipid) but not in DDM micelles containing POPS (net negative) (Fig. 1e). This effect on β_2_AR mediated turnover was significant, even though the lipid:DDM molar ratio was only 1:4. The lipid environment (POPS vs. POPE) did not affect basal GTP turnover by G_i3_ (Supplementary Fig. 3). Taken together, these results indicate that the charge property of phospholipids regulates G_i_ activation by β_2_AR.

### Ca^2+^ promotes β_2_AR-G_i3_ coupling in negatively charged phospholipids

Ca^2+^, a ubiquitous second messenger, plays an important role in cardiac myocytes; Ca^2+^ waves, magnified by G_s_ activation, drive the cardiac myocyte contraction machinery. Recently Ca^2+^ was reported to regulate T cell receptor activation by modulating the charge property of lipids^30^. Given that Ca^2+^ interaction with negatively charged phospholipids screens the negative charge, we tested whether Ca^2+^ improves mB-β_2_AR- G_i3_ coupling efficiency in negatively charged DDM-POPS. Indeed, Ca^2+^ improved coupling efficiency in DDM-POPS micelles (Fig. 2a), and this effect required POPS (Fig. 2b). Moreover, Ca^2+^ had little effect on mB-β_2_AR-G_s_ interaction, implicating differences in G_s_ and G_i3_ surface charge.

**Figure 2.**
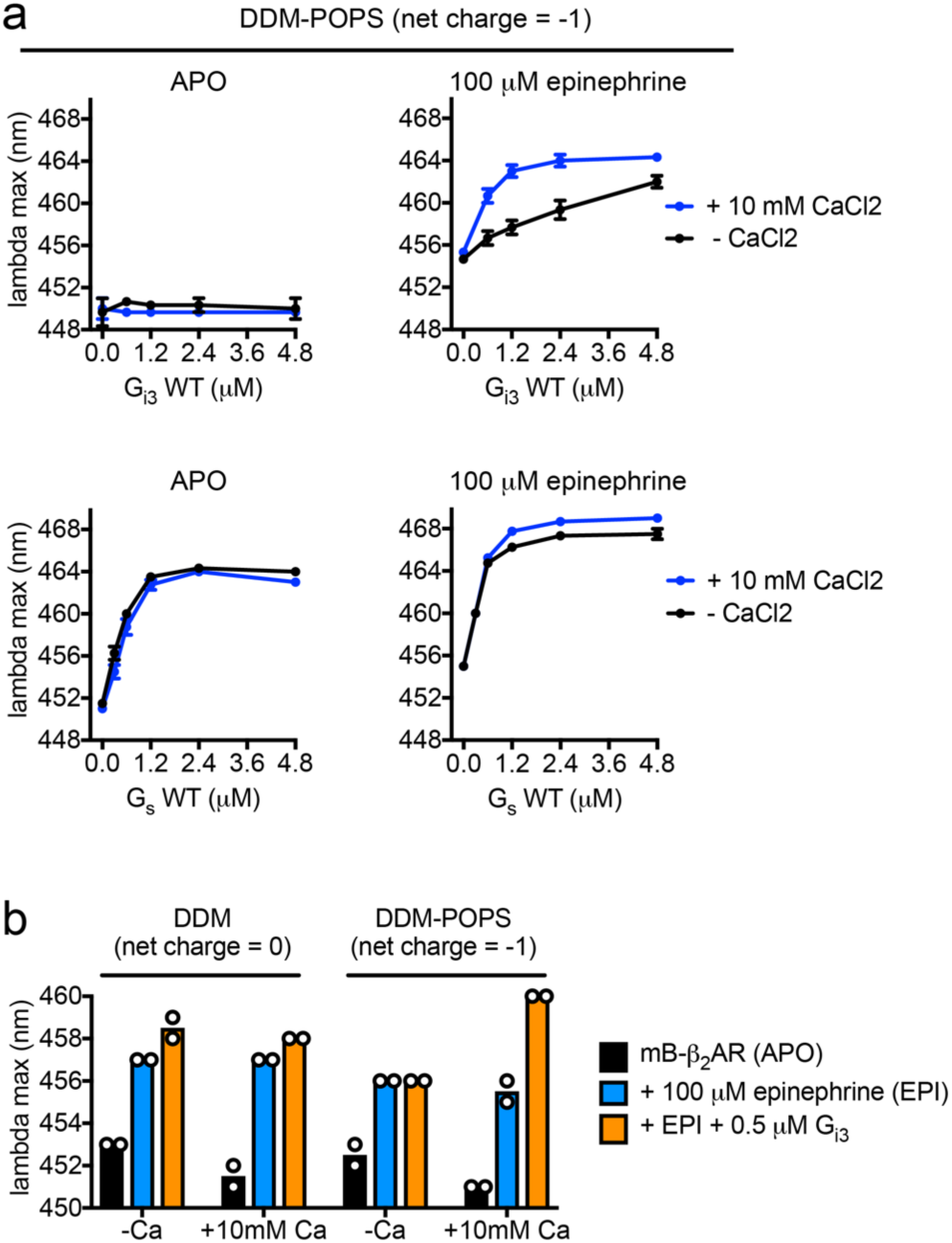
The effect of Ca^2+^ on β_2_AR-G_i3_ and β_2_AR-G_s_ interaction in micelles containing negatively charged phosphatidylserine (POPS) **a.**The effect of 10 mM CaCl2 on mB-β_2_AR-G protein interaction (G_i3_ and G_s_) was examined in micelles containing 4:1 DDM:POPS. Data were collected in the absence (left) and presence (right) of epinephrine. mB-β_2_AR concentration is 250 nM. Data are mean +/- s.e.m of three independent experiments. **b.**The effect of 10 mM CaCl2 on mB-β_2_AR-G_i3_ interaction in DDM micelles +/- POPS (i.e. 100% DDM vs. 3:2 DDM:POPS). mB-β_2_AR concentration is 250 nM. Data are mean of two independent experiments.

### Ca^2+^ interacts with the amino terminal helix of G_i3_

Next, we sought to determine the mechanism by which Ca^2+^-POPS interactions increase mB-β_2_AR coupling to G_i3_ but not to G_s_. Given that the amino terminal helix (αN) of G protein is adjacent to the membrane when coupled to the β_2_AR^27^, and polybasic residues on G_s_ αN are known to facilitate membrane interaction^31^, we looked for a possible selectivity determinant within αN. Given αN of G_s_ and G_i_ are differentially charged (Fig. 3a), we first replaced αN of G_s_ with αN of G_i3_, creating a G_i3_-G_s_ chimera (Fig. 3b).

**Figure 3.**
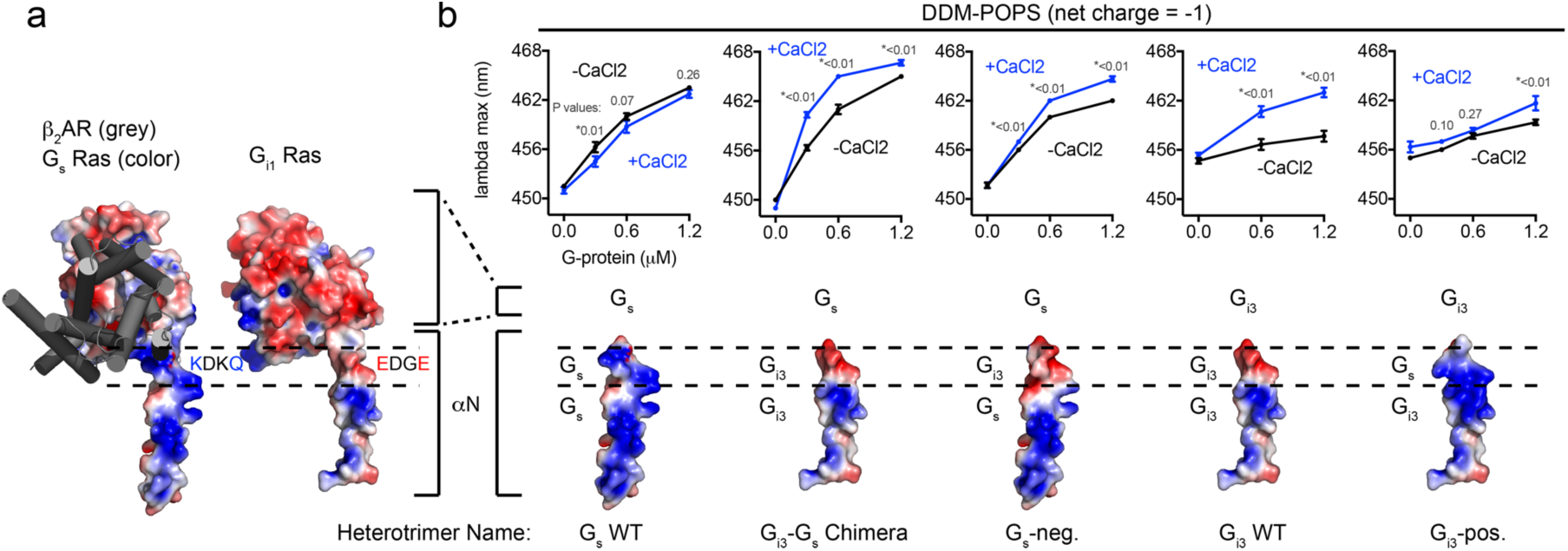
The effect of Ca^2+^ on mB-β_2_AR interaction with G protein charge mutants. a.Models of the membrane-facing surfaces of the Ras domains of G_s_ (PDB: 3SN6) and G_i1_ (1GP2). The membrane-facing surface of G_i1_ was modeled by superimposing the Ras domain of Gi1 onto the structure of G_s_ in complex with β_2_AR (PDB: 3SN6). Red and blue signify negative and positive charge, respectively. β_2_AR is shown in grey. Dashed lines highlight the region in *α*N where charge differs: The sequence is KDKQ in WT G_s_ vs. EDGE in WT G_i1_ (and in G_i2_, G_i3_). mB-β_2_AR-G protein dose-response curves +/- 10 mM CaCl2. Data were generated with the G protein mutant depicted below the curves: mutations were made in *α*N and corresponding electrostatic models are shown. Epinephrine was not included in experiments titrating G_s_ WT, G_i3_-G_s_ Chimera, or with G_s_-neg. to enhance the effect of CaCl2. Epinephrine (100 µM) was included in experiments titrating G_i3_ WT and G_i3_- pos. mB-β_2_AR concentration is 250 nM. Data are mean +/- s.e.m of three independent experiments. Statistical significance was determined by using a two-sided Student’s *t*-test. Test assesses the effect of CaCl2 at each G protein concentration.

While Ca^2+^ does not promote mB-β_2_AR coupling to WT Gs (Fig. 3b), it did promote mB- β_2_AR coupling to the G_i3_-G_s_ chimera (Fig. 3b). Next, we compared the membrane-facing charge of G_s_ WT αN and G_i3_ WT αN. Structural analysis revealed that charge differed at the C terminal end of αN: G_i_ harbors a negatively charged motif (EDGE) at the position where G_s_ harbors a positively charged motif (KDKQ) (Fig 3a). To examine whether this motif dictates a differential response to Ca^2+^, we constructed a G_s_ mutant (“G_s_-neg.”) containing the negatively charged motif of G_i3_ (KDKQ?EDGE). Ca^2+^ increased mB-β_2_AR interaction with this mutant (Fig 3b), suggesting the EDGE motif is responsible for the effect of Ca^2+^ on G_i3_ αN. Taken together, our results imply that Ca^2+^ coordinates an interaction between the negatively charged EDGE motif on αN of G_i3_ and the headgroup of POPS. In the absence of Ca^2+^, like-charge repulsion decreases mB-β_2_AR coupling to G_i3_.

We also constructed a G_i3_ mutant (“G_i3_-pos.”) containing the positively charged motif of G_s_ (EDGE→KDKQ). The mutations only partially removed the effect of Ca^2+^ (Fig. 3b), indicating the effect of Ca^2+^ on G_i3_ extends beyond an effect on αN (see discussion).

### Bilayer charge is a tunable modulator of the G protein subtype selectivity

To examine the effects of phospholipids in a more native environment, we reconstituted mB-β_2_AR into nanodisc bilayers and purified the nanodiscs to homogeneity using size-exclusion chromatography (Supplementary Fig. 4).

First, we compared the influence of lipid composition in the absence of Ca^2+^. Negatively charged bilayers (DOPG and DOPS bilayers) red-shifted the emission spectra of mB- β_2_AR alone, suggesting negatively charged bilayers stabilize mB-β_2_AR in an active conformation, as has been previously reported^24^. While negatively charged bilayers increased mB-β_2_AR coupling to G_s_ (Fig. 4), negatively charged bilayers (especially DOPS bilayers) decreased mB-β_2_AR coupling to G_i3_ (Fig. 4).

**Figure 4.**
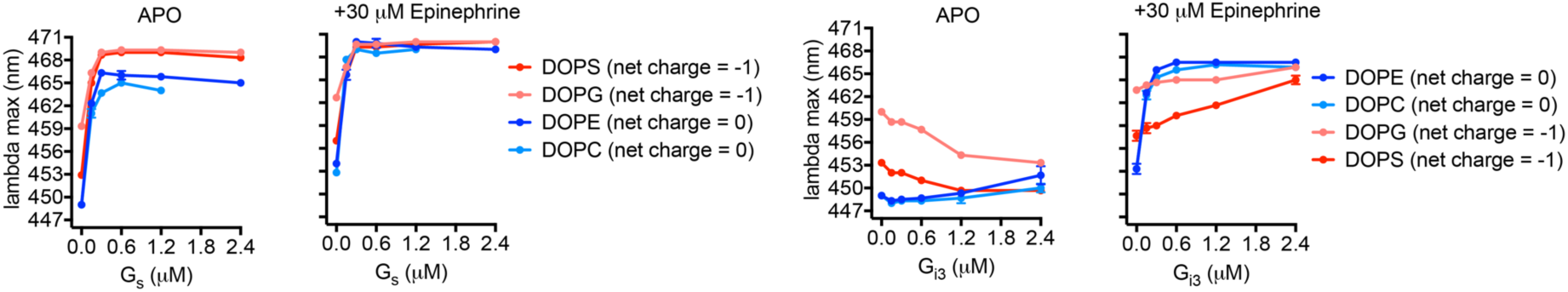
Effect of phospholipid bilayer charge on mB-β_2_AR-G_i3_ and mB-β_2_AR-G_s_ interaction. The effect of G_s_ (left) and G_i3_ (right) concentration on mB-β_2_AR fluorescence was examined in nanodisc bilayers of varying phospholipid composition (DOPE, DOPC, DOPG, or DOPS). Net charge of phospholipid indicated in parentheses. mB-β_2_AR concentration is 100 nM; maximum stoichiometry is 24:1 (G protein:mB-β_2_AR). Data are mean +/- s.e.m of three independent experiments.

In fact, in negatively charged bilayers without epinephrine, G_i3_ unexpectedly *blue-shifted* the emission spectra of mB-β_2_AR. While this may indicate that G_i3_ stabilizes the β_2_AR in an inactive conformation in negatively charged lipids, it may represent a non-specific interaction of inactive G_i3_ with the β_2_AR or the lipid bilayer.

Next we examined the effect of Ca^2+^ and Mg^2+^. In the absence of G_i3_, both Ca^2+^ and Mg^2+^ reversed the active-state stabilizing effect of negatively charged DOPS and DOPG bilayers (Fig. 5a). In contrast, Ca^2+^ and Mg^2+^ increased mB-β_2_AR coupling to G_i3_ in negatively charged DOPS bilayers, but only Ca^2+^ was efficacious at concentrations below 1 mM (Fig. 5a). Ca^2+^ similarly affected mB-β_2_AR-G_i3_ interaction in negatively charged DOPG bilayers (Fig. 5a), but the magnitude of the effect in DOPG bilayers was less than observed in DOPS bilayers due to the higher baseline effect of DOPG on β_2_AR conformation.

**Figure 5.**
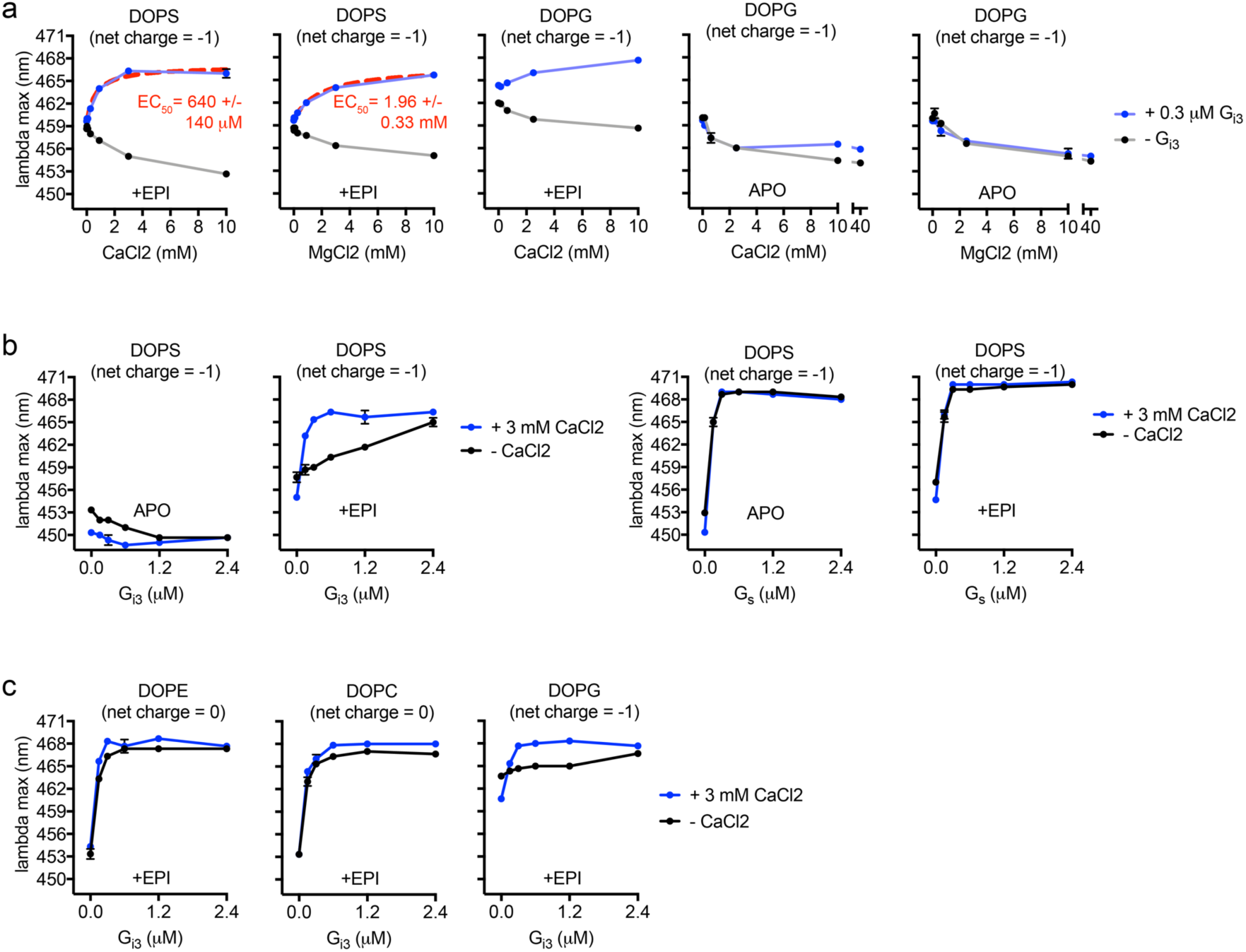
Effect of Ca^2+^ and Mg^2+^ on mB-β_2_AR-G_i3_ and mB-β_2_AR-G_s_ interaction in nanodisc bilayers of varying charge. a. The effect of CaCl2 and MgCl2 concentration on mB-β_2_AR fluorescence in DOPS and DOPG nanodisc bilayers was examined in the presence and absence of G_i3_. Epinephrine was included (30 µM) or omitted (APO). EC_50_ is mean +/- s.e.m. b. The effect of G protein concentration (G_i3_ and G_s_) on mB-β_2_AR fluorescence +/- 3 mM CaCl2 was examined in DOPS nanodiscs in the presence of epinephrine (30 µM) or in its absence (APO). The effect of G_i3_ on mB-β_2_AR fluorescence +/- 3 mM CaCl2 was examined in DOPE, DOPC, and DOPG nanodisc bilayers in the presence of 30 µM epinephrine (EPI). mB-β_2_AR concentration is 100 nM. Net charge of phospholipid molecule indicated in parentheses. Data are mean +/- s.e.m of three independent experiments.

We compared the effect of Ca^2+^ on mB-β_2_AR interactions with G_s_ and G_i3_ in DOPS bilayers. As observed in micelles, Ca^2+^ increased mB-β_2_AR coupling to G_i3_ but not to Gs (Fig. 5b). Ca^2+^ also improved mB-β_2_AR-G_i3_ coupling efficiency in negatively charged DOPG bilayers (Fig. 5c). Only a minor effect of Ca^2+^ was observed in neutral DOPE and DOPC bilayers (Fig. 5c & Supplementary Fig. 5), which could be attributable to weaker Ca^2+^/DOPE and Ca^2+^/DOPC interactions that have been reported^32^. Taken together, our results strongly suggest that Ca^2+^ facilitates mB-β_2_AR-G_i3_ interaction but not mB-β_2_AR- G_s_ interaction. Moreover, these observations provide biochemical proof-of-concept that divalent cation/membrane interactions can increase G_i_ coupling to the β_2_AR.

## Discussion

We observed that local membrane charge regulates β_2_AR-G protein interaction. Negatively charged membrane promotes β_2_AR-G_s_ coupling and suppresses β_2_AR-G_i3_ coupling. However, G_s_ bias is reduced in neutral membrane and in negatively charged membrane in the presence of divalent cations (see model in Fig. 6).

**Figure 6.**
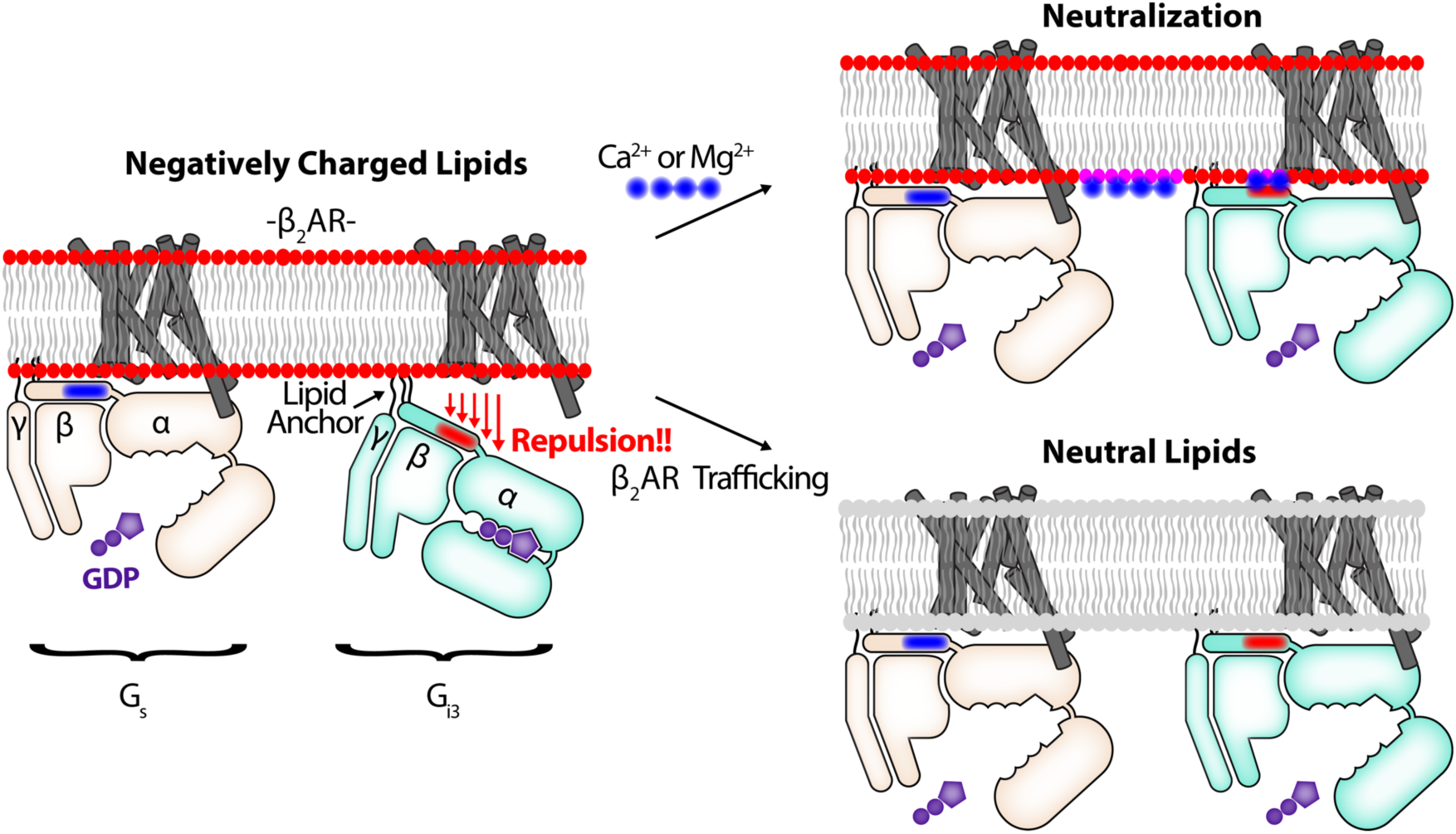
Membrane charge is a tunable modulator of β_2_AR-G protein interaction. Models depict epinephrine-bound β_2_AR. **Left:** In negatively charged lipids, β_2_AR-G_s_ coupling is efficient, but β_2_AR-G_i3_ coupling is relatively inefficient, in part because β_2_AR-G_i3_ attraction is countered by membrane-G_i3_ repulsion. Specifically, negatively charged lipids repel the negatively charged EDGE motif found on the amino terminal helix of G_i3_ (shown in red), a region that is positively charged in G_s_ (shown in blue). **Right:** Two mechanisms that neutralize membrane charge facilitate β_2_AR coupling to G_i3_. These mechanisms may play a role in G_s_-to-G_i_ switching in cardiac myocytes. Top right: Ca^2+^ and Mg^2+^ stabilize a like-charge interaction between the membrane and the EDGE motif. (Note that the effect of Ca2+ and Mg^2+^ may extend beyond an effect on *α*N positioning). Bottom right: Epinephrine stimulated β_2_AR traffics to membrane without negatively charged lipids.

We have begun to explore the mechanism by which Ca^2+^ increases mB-β_2_AR coupling to G_i3_ in PS phospholipids. Although G proteins are membrane tethered via lipidation, the lipid anchor of G_i3_ is not sufficient for optimal interaction with β_2_AR in negatively charged bilayers, possibly due to repulsion of the carboxyl terminal end of the αN helix. We propose that Ca^2+^ helps orient the carboxyl terminal end of the αN helix of G_i3_ near the membrane, thereby facilitating β_2_AR-G_i3_ interactions. More specifically, we propose that Ca^2+^ facilitates the interaction of PS with the negatively charged EDGE motif on the αN helix of G_i3_. Ca^2+^ may stabilize the PS-EDGE interaction by coordinating a like-charge interaction between the carboxylate groups on G_i3_ and the carboxylate group on PS (not present on PG) (refer to structures in Fig 1b). Alternatively, Ca^2+^ might coordinate a like-charge interaction between the carboxylate groups on G_i3_ and the phosphate group present on PS and PG, or Ca^2+^ might coordinate an intramolecular interaction between the phosphate group and the carboxylate group on PS, “freeing” the amino group (NH3+) on PS to interact with the carboxylate groups on G_i3_. Notably, the amino terminal helix (αN) of G_i1_, G_i2_, and G_i3_ are similarly charged, and all three sequences contain the EDGE motif.

We have previously shown that negatively charged lipids, particularly PG, stabilize the β_2_AR in an active-like conformation as revealed by changes in mB-β_2_AR fluorescence and an increased affinity for agonists^24^. These effects are likely due to interactions between the lipids and positively charged amino acids on the β_2_AR. Here we observed that the effect of DOPG and DOPS on mB-β_2_AR can be reversed by both Ca^2+^ and Mg^2+^ (Fig. 5a). Yet, these divalent cations do not appear to reduce coupling to G_s_.

β_2_AR signals from caveolin-rich rafts^33,34^ within T-tubules^35^. While β_2_AR preferentially interacts with PG in insect cell membrane^24^, the phospholipid composition immediately adjacent to β_2_AR in T-tubules, and how it changes during β_2_AR trafficking, is currently unknown. Net-neutral PC and PE are the major phospholipids in T-Tubules^36-38^.However, negatively charged PS is enriched in T-tubules relative to other membrane fractions (7.5-12.3% of total phospholipid)^36-40^. While cytosolic Ca^2+^ concentrations are typically less than 1 mM^41^, concentrations of Ca^2+^ in the mM range may be observed in cardiac myocytes (discussed below).

Investigators have long speculated about the functional role of Ca^2+^ in the cleft between T-tubule membrane (where β_2_AR is localized) and juxtaposed sarcoplasmic reticulum (SR)^42-44^. During each action potential, extracellular Ca^2+^ flows into the cleft through L-type Ca^2+^ channels (LTCCs) on the plasma membrane and through ryanodine receptors (RyRs) on the sarcoplasmic reticulum^41^. Cleft Ca^2+^ concentrations spark to > 100 µM in the absence of epinephrine and >1 mM^45,46^ following epinephrine stimulation, a consequence of G_s_ activation. Computational models show that negatively charged phospholipids buffer approximately half the Ca^2+^ released into the cleft^45^, and experiments have shown that 80% of inner-leaflet bound Ca^2+^ is bound to negatively-charged phospholipids^47^. Additionally, biochemical investigations show that Ca^2+^ can cluster negatively charged PS^48^ and PIP2^49,50^ lipids.

β_1_AR and β_2_AR signaling through G_s_ alters calcium handling in the cardiac myocyte, and increases the magnitude of Ca^2+^ currents and Ca^2+^ transients, which stimulate cardiac contraction^41,51^. However, elevated Ca^2+^ concentrations also activate the Ca^2+^/calmodulin-dependent protein kinase II (CaMKII), which is implicated in structural remodeling that ultimately results in cardiac dysfunction^52-56^. Several lines of evidence suggest β_2_AR-G_i_ signaling keeps β_2_AR-G_s_ signaling in check via negative feedback: β_2_AR-G_i_ signaling occurs minutes after β_2_AR-G_s_ signaling^3^, β_2_AR-G_i_ signaling suppresses changes in calcium handling^51,57^, and β_2_AR-G_i_ signaling is anti-apoptotic^7,8^. While the mechanism that triggers β_2_AR-G_i_ signaling is unknown, our biochemical observations suggest Ca^2+^ concentrations could directly regulate β_2_AR coupling to G_i_. It is notable that overexpression of the Ca^2+^/sodium exchanger facilitates β_2_AR-G_i_ suppression of β_1_AR-G_s_ signaling^58^, and overexpression has been cited to increase the inward LTCC Ca^2+^ current^59^.

It is also notable that intracellular Ca^2^+^60,61^ and Mg^2^+^61^ concentrations rise during ischemia and rise even higher during reperfusion. Whether the rising concentrations affect the G protein subtype specificity of β_2_AR is unknown. However, laboratory-controlled bouts of ischemia and reperfusion have been cited to evoke therapeutic β_2_AR- G_i_ signaling, i.e. reduce necrosis during a heart attack^9^.

Whether negatively charged phospholipids affect G_i_ interaction with other G_i_-coupled GPCRs is not known. The observation that Ca^2+^ sensing receptor (CaSR) switches from Gq to G_i_ after cytosolic Ca^2+^ increases^62^ is potentially relevant to our findings.

In conclusion, we show that local membrane charge differentially modulates β_2_AR interaction with competing G protein subtypes (G_s_ and G_i_). This discovery expands our knowledge of mechanisms that regulate the G protein coupling selectivity of GPCRs.

## Online Methods

### Heterotrimeric G protein and β_2_AR purification

All G proteins were heterotrimeric G proteins. G_s_ heterotrimer (WT G*α*s short, his6-3C-*β*1, WT γ2) and G_i_ heterotrimer (WT G*α*i1- 3, his6-3C-*β*1, WT γ2) and all G protein mutants were expressed and purified as previously described^63^. The G_i3_-G_s_ chimera was constructed by replacing residues 1-38 of WT G*α*s with residues 1-31 of WT G*α*i3. The G_s_-neg. mutant was constructed by replacing the sequence EDGE (residues 25-28) of WT G*α*s with the sequence KDKQ. The G_i3_-pos. mutant was constructed by replacing the sequence KDKQ (residues 32-35) of WT G*α*i3 to the sequence EDGE. The β_2_AR construct was PN1. The construct, expression, purification, and monobromobimane labeling have been described^24^.

Labeling efficiency was ∼80-100%, as determined by spectroscopic analysis. Purified β_2_AR and G protein were dephosphorylated using Lambda Protein Phosphatase (Lambda PP, New England BioLabs, NEB). Where indicated, β_2_AR was phosphorylated with Protein Kinase A (PKA, NEB) in purification buffer supplemented with 0.1 mM EDTA and 20 µM ATP. (Subsequently, EDTA/ATP were removed by dialysis). Phosphorylation was assessed using the Pro-Q Diamond Phosphoprotein Gel Stain (ThermoFisher Scientific), per the manufacturer’s instructions.

### Micelle Composition

n-dodecyl-*β*-D-maltopyranoside (DDM), Cholesteryl hemisuccinate (CHS), and 1-palmitoyl-2-oleoyl-(PE,PC,PG,PS) lipids (Avanti Polar Lipids) were mixed in the indicated ratios and solubilized in chloroform. Chloroform was evaporated, and films were re-suspended in 20 mM HEPES (pH 7.4), 100 mM NaCl.

### Fluorescence Spectroscopy

In experiments examining β_2_AR in micelles, mB-β_2_AR was pre-incubated (30 min room temperature) in micelle stock prior to dilution with other reaction components. Buffer (containing 100 mM NaCl, +/- ligand, +/- CaCl2 or MgCl2), and G protein were sequentially included. Mixtures were incubated 2.5 – 3.0 h at room temperature. Final mB-β_2_AR concentration was 100-500 nM. Emission spectra were read at 22 degrees Celsius using Flurolog-3 or Spex FluoroMax-3 spectrofluorometers (Horiba Jobin Yvon Inc.). (Bandpass = 4 nm; Excitation = 370 nm; Emission = 420-500 nm.) Raw S1c/R1c spectra were smoothed using Prism (GraphPad Software) (n=15 neighbors, 2^nd^ order polynomial). Lambda max is defined as the wavelength at which fluorescence emission is maximum. To determine the EC_50_, curves were fit to the “agonist vs. response” model in Prism 7.0d software.

### GTP Turnover

Samples were prepared as they were for fluorescence spectroscopy. Following the incubation with G protein, 1 µM GTP/5 µM GDP mixtures were added. Reactions contained: 20 mM HEPES (pH 7.4), 100 mM NaCl, 0.25 µM G_i3_, 0.5 or 1.0 µM β_2_AR (per legend), 200 µM epinephrine, and 4:1 DDM:Lipid. 12 minutes after GTP was added, free GTP was assessed using the GTPase-Glo assay (Promega), which reports free GTP concentration using a luminescence readout. Luminescence was detected using a SpectraMax Paradigm plate reader equipped with a TUNE SpectraMax detection cartridge (Molecular Devices). Background luminescence was subtracted from experimental reactions.

### Statistics

Two-sided parametric paired student’s *t*-tests were performed using Graphpad Prism7.0d. When comparing lambda max +/- CaCl2, *t*-test calculation assumed all data within a panel were sampled from populations with the same scatter.

### Electrostatic Modeling

Structural views and mutant models were generated using PyMOL (Schrödinger, LLC). We selected rotomer positions that most closely matched those seen in PDB 3SN6 (for “G_i3_-pos.” model) and PDB 1GP2 (for “G_s_-neg.” model). Continuum electrostatics models were calculated using the APBS^64^ plugin (MG Lerner,University of Michigan, Ann Arbor) for PyMOL. Atomic charge and radii were calculated using the online PDB2PQR server^65^ (pH 7.4, PARSE force field, hydrogen bond optimization, clash avoidance).

### Nanodisc Reagents

1,2-dioleoyl-(PE,PC,PG,PS) lipids (Avanti Polar Lipids) were used because of their low phase transition temperature. MSP1E3D1 (Addgene #20066) was expressed and purified as described^66^.

### Nanodisc Reconstitution

Reconstitution was performed as described^24^ with the following modifications: Nanodiscs were formed with one lipid type (i.e. 100% DOPS, DOPG, DOPE, or DOPC). The Lipid-to-MSP1E3D1 ratio was 35:1. The MSP1E3D1-to- mB-β_2_AR ratio was 1:10. Empty nanodiscs were separated from nanodiscs containing mB-β_2_AR using M1-anti-FLAG immunoaffinity chromatography in the presence of 2 mM CaCl2, which captures the FLAG-tagged PN1 mB-β_2_AR construct. Eluate was incubated with 5 mM EDTA > 1.5 h at 4 degrees Celsius to remove divalent cations. Subsequently,samples were injected into a Superdex 200 10/300GL size-exclusion column (GE Healthcare) and the main peak was harvested. The concentration of nanodisc β_2_AR was approximated by SDS-PAGE, using detergent purified β_2_AR of known concentration as a reference.

## Acknowledgements

We thank Betsy White for assistance with G protein and β_2_AR expression. This work was supported by National Institutes of Health grants R01NS028471 and R01GM083118 (B.K.K.). B.K.K. is supported by the Chan Zuckerberg Biohub. D.H. was supported by the German Academic Exchange Service (DAAD). M.M. was supported by the American Heart Association Postdoctoral fellowship (17POST33410958).

## Author contributions

M.J.S designed, performed, and interpreted the research, and wrote the manuscript.B.K.K. championed the investigation, advised on the project, and edited the manuscript.M.J.S performed the purification, labeling, nanodisc reconstitution, and all of the experiments, and S.M., D.H., M.M., and Y.D. provided valuable technical assistance as described: S.M. advised on cloning and provided G protein for pilot experiments. D.H. advised on G protein purification and the GTP turnover assay. M.M. advised on nanodisc reconstitution and provided MSP1E3D1 protein. Y.D. collaborated on pilot experiments not included.

